# Enhancing c-MYC degradation via 20S proteasome activation induces *in vivo* anti-tumor efficacy

**DOI:** 10.1101/2020.08.24.265470

**Authors:** Evert Njomen, Theresa A. Lansdell, Allison Vanecek, Vanessa Benham, Matt P. Bernard, Ya-Ting Yang, Peter Z. Schall, Daniel Isaac, Omar Alkharabsheh, Anas Al-Janadi, Matthew B. Giletto, Edmund Ellsworth, Catherine Taylor, Terence Tang, Sarah Lau, Marc Bailie, Jamie J. Bernard, Vilma Yuzbasiyan-Gurkan, Jetze J. Tepe

**Affiliations:** Department of Chemistry, Michigan State University, East Lansing, Michigan 48824, United States; Department of Pharmacology & Toxicology, Michigan State University, East Lansing, Michigan 48824, United States; Comparative Medicine and Integrative Biology Program, Michigan State University, East Lansing, Michigan 48824, United States; Breslin Cancer Center and Clinical Center, Michigan State University, East Lansing, Michigan 48824, United States; Department of Microbiology and Molecular Genetics, Michigan State University, East Lansing, Michigan 48824, United States; Department of Small Animal Clinical Sciences, Michigan State University, East Lansing, Michigan 48824, United States; Department of Biology, University of Waterloo, ON, Canada

**Author notes:** Correspondence (JJT), (VYG).

## Abstract

Enhancing proteasome activity is a potential new therapeutic strategy to prevent the accumulation of aberrant high levels of protein that drive the pathogenesis of many diseases. Herein, we examine the use of small molecules to activate the 20S proteasome to reduce aberrant signaling by the undruggable oncoprotein c-MYC, to treat c-MYC driven oncogenesis. Overexpression of c-MYC is found in more than 50% of all human cancer but remains undruggable because of its highly dynamic intrinsically disordered 3-D conformation, which renders traditional therapeutic strategies largely ineffective. We demonstrate herein that small molecule activation of the 20S proteasome targets dysregulated intrinsically disordered proteins (IDPs), including c-MYC, and reduces cancer growth *in vitro* and *in vivo* models of multiple myeloma, and is even effective in bortezomib resistant cells and unresponsive patient samples. Genomic analysis of various cancer pathways showed that proteasome activation results in downregulation of many c-MYC target genes. Moreover, proteasome enhancement was well tolerated in mice and dogs. These data support the therapeutic potential of 20S proteasome activation in targeting IDP-driven proteotoxic disorders, including cancer, and demonstrate that this new therapeutic strategy is well tolerated *in vivo*.

## INTRODUCTION

The induction of protein degradation as a new therapeutic strategy has gained significant excitement and validation.^1^ The protease responsible for targeted protein degradation is the human proteasome, consisting of an equilibrium of various isoforms, including the 26S and 20S isoforms. Currently, the protein degradation approach utilizes a proteolysis targeting chimera (PROTAC) strategy that relies on the ubiquitin-dependent degradation of a target protein by the 26S proteasome.^2^ The 20S isoform of the proteasome targets intrinsically disordered proteins (IDPs) directly via a ubiquitin-independent mechanism.^3-5^ Enhancing the proteolytic activity of only the 20S isoform of the human proteasome represents a new strategy to reduce aberrant signaling of dysregulated IDPs.^6-11^ IDPs are typically short-lived and are rapidly and unremittingly degraded by the proteasome. However, when production of a specific IDP outpaces proteasome degradation, the accumulated IDP can induce aberrant and toxic signaling events. Dysregulated IDPs are involved in the pathogenesis of many diseases including neurodegenerative disorders and cancer.^6,7,12,13^

Overexpression of the intrinsically disordered oncoprotein, c-MYC, is found more than 50% of all human cancer, with high prevalence in cancers of hematopoietic origin, such as acute lymphoblastic leukemia and multiple myeloma.^14^ In normal cells, c-MYC protein levels are kept in check through multiple post-translational modifications that alter susceptibility toward proteasomal degradation by default.^15^ Dysregulation of c-MYC clearance can induce its oncogenic transcriptional activity, resulting in the transcription of multiple tumor promoting genes.^16^ Due to its highly dynamic intrinsically disordered 3-D conformation, c-MYC has evaded pharmacological manipulation and remains one of the most sought of anticancer targets.^17^

We evaluated the use of 20S proteasome enhancement for its therapeutic potential in cell culture and *in vivo*, using the 20S proteasome enhancer, TCH-165. TCH-165 is part of a privileged class of imidazoline-based scaffolds,^18-24^ and is one of the few known molecules shown to enhances 20S mediated proteolysis nearly 10 fold (i.e. 1000%) by favoring a proteolytically active, open-gate 20S proteasome subcomplex.^25^ Our findings show that TCH-165 effectively reduces c-MYC protein levels *in vitro* and in cell culture, affects c-MYC target genes, inhibits cancer cell proliferation, and inhibits tumor growth *in vivo*. Importantly, this proteasome enhancer also effectively kills cancer cells resistant to proteasome inhibitors, including primary cells from bortezomib unresponsive MM patients. Moreover, the 20S proteasome enhancer is well tolerated *in vivo* (in mice and in dogs) at therapeutic concentrations. These studies support further exploration of 20S proteasome enhancement as a new and well-tolerated approach to treat IDP-driven disorders, including c-MYC driven oncogenesis.

## RESULTS

### TCH-165 enhances the proteolytic activity of the 20S proteasome

The 20S proteasome is a barrel-shaped multi-subunit protease with 28 subunits arranged in 4 stacked heptameric rings (***Fig. 1A***). Its inner β-rings contain 3 catalytic subunits (β5, β2 and β1) that exhibit chymotrypsin-like (CT-L), trypsin-like (Tryp-L) and caspase-like (Casp-L) activities.^26^ Its outer α-rings guard entrance into the proteolytic core, via an allosterically-controlled gate-opening/closing mechanism, which is required for substrate access to the proteolytic sites.^27^ In a previous mechanistic study, TCH-165 increased substrate accessibility to the 20S catalytic chamber through 20S gate opening (***Fig. 1A***).^25^ TCH-165 and other 20S activators identified in our lab were reported to only enhance the degradation of endogenous 20S substrates.^25,28^ Structured proteins normally targeted by the 26S-ubiquitin-dependent proteasome system were not affected. To determine the Active Concentration (AC) at which TCH-165 doubles (200%) 20S activity (referred to hereafter as AC_200_), and the maximum fold enhancement of 20S activity (referred to hereafter as Fold^M^), the hydrolysis of canonical proteasome peptide substrates (***Fig. 1B***) were measured in the presence of TCH-165. The AC_200_ for the hydrolysis of the CT-L substrate, Suc-LLVY-AMC was 1.5μM, with Fold^M^ of 810%. The AC_200_ and Fold^M^ for the Tryp-L and Casp-L sites were also determined to be 2.7μM (Fold^M^ 500%) and 1.2μM (Fold^M^ 1290%), respectively.

**Figure 1:**
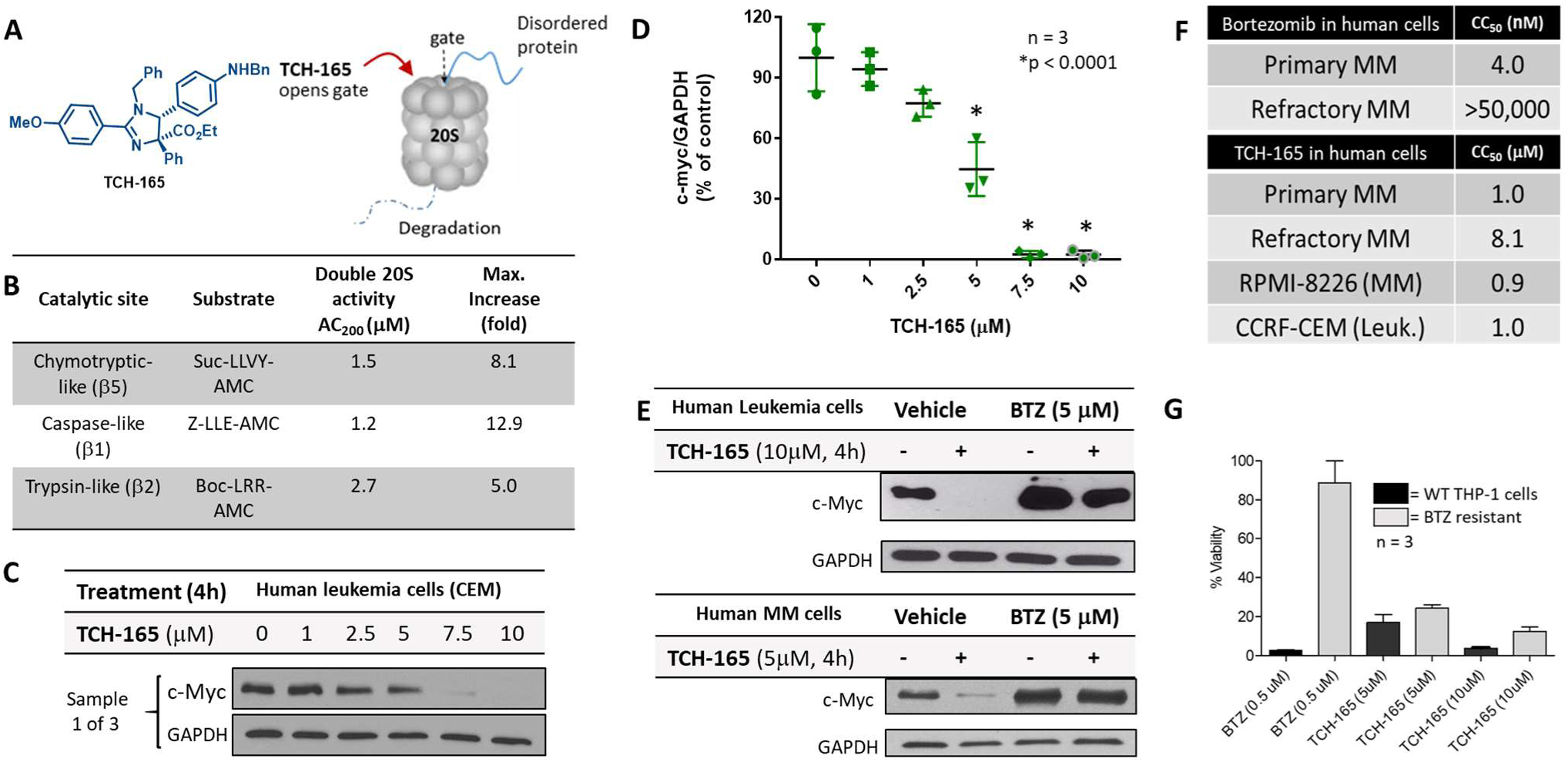
Enhanced degradation of c-MYC and susceptibility of cancer cells to the proteasome activator, TCH-165. **(A)** Structure of TCH-165 and its proposed model of α-ring binding allowing the gate to open for easy access of disordered proteins to the internal proteolytic sites. **(B)** AC_200_ values of TCH-165 and maximum fold enhancement of 20S activities for each of the three catalytic sites. **(C)** Sample 1 of 3 of immunoblot of cell lysates showing concentration dependent reduction of c-MYC in human leukemia cells upon 4h treatment with various concentrations of TCH-165 (Fig. 1S. for additional representative samples, n=3) **(D)** Statistical analysis of c-MYC reduction from Western blots (Fig. S1, n=3) using one-way ANOVA, with Sidak’s multiple comparisons test. **(E)** Abrogation of TCH-165 induced c-MYC degradation following BTZ-induced proteasome inhibition in both CEM and RPMI-8226 cells. **(F)** Cytotoxicity of TCH-165 in primary human MM cells of newly diagnosed patient (CC_50_ 1.0μM; 95% CI 0.60-1.51μM), relapsed patient (CC_50_ 8.1μM; 95% CI 7.08-9.03μM), as well as human RPMI-8226 (CC_50_ 1.0 μM; 95% CI 0.75-1.15μM) and CCRF-CEM (CC_50_ 0.9 μM; 95% CI 0.79-1.19μM) cells after 72h treatment. **(G)** Viability of wild-type and BTZ-resistant human cute monocytic leukemic THP-1 cells exposed to TCH-165 (72h) at 5μM and 10μM (n=3). 95% CI = 95% confidence interval.

### c-MYC degradation by the 20S proteasome is enhanced by TCH-165

First, we confirmed that the IDP c-MYC is targeted for degradation by the 20S proteasome. For these studies, increasing concentrations of purified 20S proteasome were incubated with c-MYC. Western blot analysis of the *in vitro* system clearly showed a decrease in the amount of c-MYC with increasing concentration of 20S proteasome (***Fig. 1S***). These results indicate that c-MYC is a target of the 20S proteasome.

Considering the critical role of c-MYC in driving tumor growth and relapse in hematological cancers, the effects of TCH-165 on the protein levels of c-MYC were evaluated in two cell lines; CCRF-CEM (human T-cell acute lymphoblastic leukemia) and RPMI-8226 (B-lymphocytes; multiple myeloma) cell lines.^29,30^ For these assays, CCRF-CEM or RPMI-8226 were treated with different concentrations of TCH-165 or with single concentration, with or without the proteasome inhibitor, bortezomib (BTZ) or 4h. BTZ was used here as a control to evaluate the effects of TCH-165 on c-MYC degradation, if proteasome activity was abrogated. As illustrated in ***Fig. 1C*** and ***1D***, TCH-165 showed a significant (n=3, \**p*=0.0001) concentration-dependent reduction in the cellular level of c-MYC (***see also supplemental data Fig. S2***). Importantly, TCH-165-induced clearance of c-MYC in both CCRF-CEM (***Fig. 1E***, top panel) and RPMI-8826 (***Fig. 1E***, bottom panel) cells were blocked by the proteasome inhibitor, bortezomib (BTZ). These observations implicated the proteasome as the likely target for the effect of TCH-165 on c-MYC. Under the same treatment conditions (5μM, 4h), the mRNA levels of c-MYC were not affected in RPMI-8226 cells (***Fig. 3***). However, in CEM cells, MYC mRNA was significantly reduced at higher concentration of TCH-165 (***Fig. 3***, 10μM, log2 > −0.5).

**Fig. 2:**
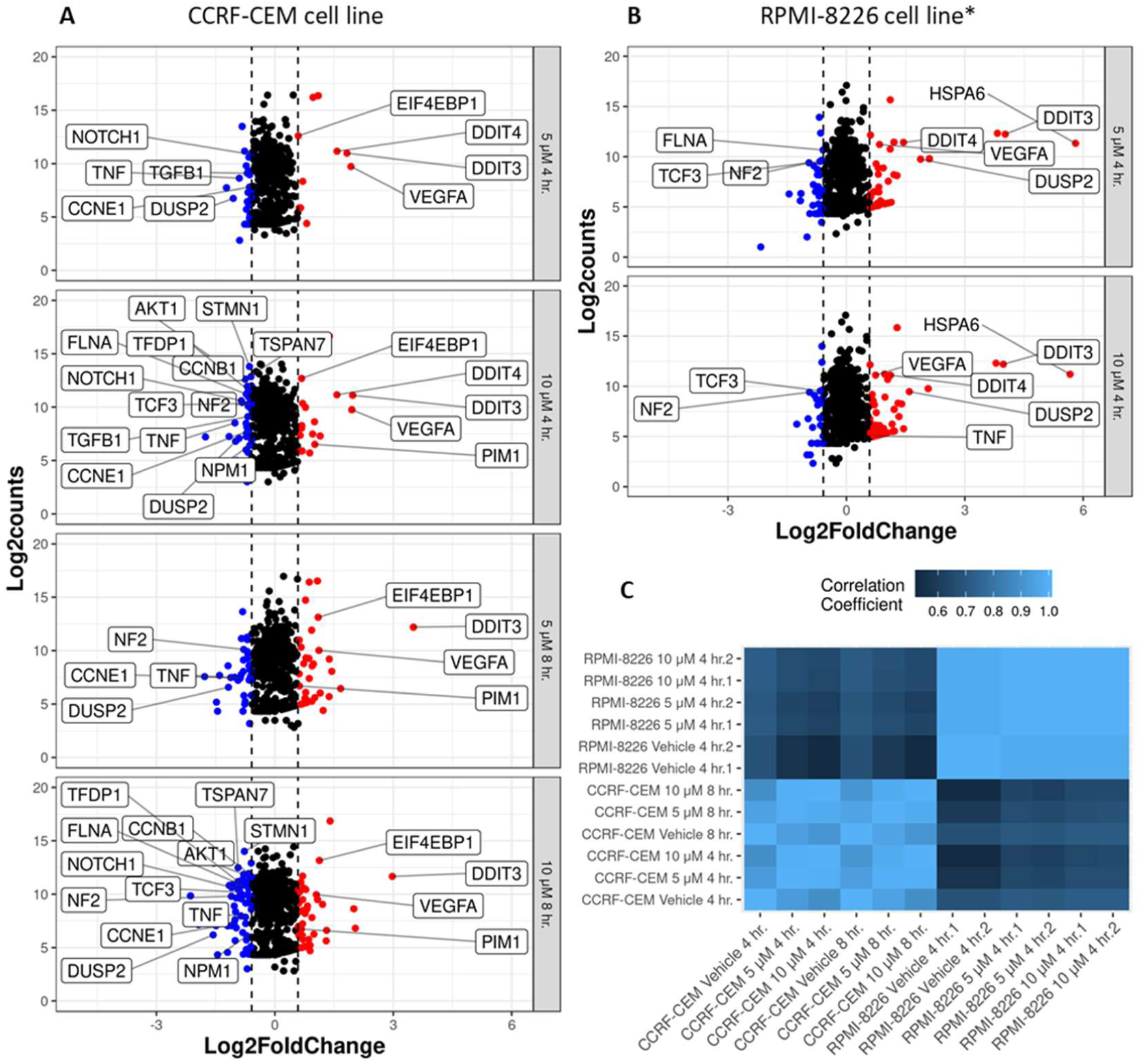
Gene Expression Profiles of CCRF-CEM and RPMI-8226 cells following TCH-165 treatment. **A-B**. Volcano scatter plots of genes in cancer pathways analyzed by the PAN CANCER™ multiplexed transcript detection assay. Dotted vertical lines indicate 1.5 fold change in normalized gene expression, with over-expressed genes in red and under-expressed genes in blue. Genes identified within boxes reflect targets of MYC identified through IPA. HSPA6, overexpressed upon treatment with TCH165 in the RPMI-826 cell line, is indicated by name. **Panel A**: Gene expression in CCRF-CEM cells treated for 4 or 8h with 5 or 10μM TCH 165 versus vehicle control. **Panel B**. RPMI-8226 cells treated for 4h with 5 or 10μM TCH-165. **Panel C**. Heatmap of pairwise correlation analysis among all samples.

**Fig. 3:**
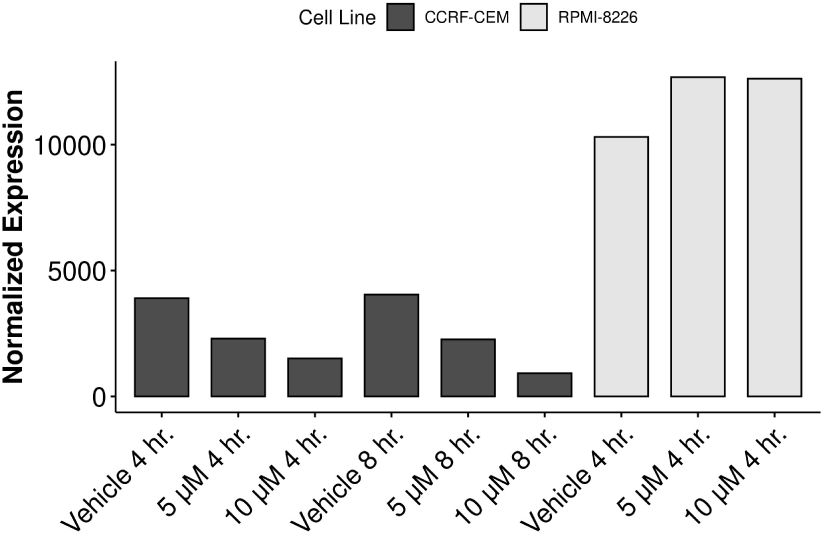
MYC expression. Normalized expression of MYC across both CCRF-CEM and RPMI-8226 cells at baseline and upon treatment with TCH-165.

### Various hematological wild-type and resistant cancer cell lines, including, patient derived primary and refractory multiple myeloma cells are vulnerable to the proteasome activator, TCH-165

To evaluate the efficacy of TCH-165, the viability of RPMI-8662, CCRF-CEM and THP-1 (acute monocytic leukemia) cells were evaluated following treatment with TCH-165. TCH-165 inhibited the proliferation of these cells with low single digit micromolar potency with a 50% cytotoxic concentration (CC_50_) of 1.0 μM (95% CI 0.75-1.15μM) in RPMI-8226 cells and 0.9 μM (95% CI 0.79-1.19μM) in CCRF-CEM cells after 72 h (***Fig. 1F***). Although bortezomib and second-generation proteasome inhibitors remain first line therapy in multiple myeloma treatment, the lack of response and relapse by a large population of patients remain a challenge.^31,32^ Accordingly, we evaluated the effect of TCH-165 and bortezomib (control) in primary cells isolated by CD138+ enrichment from newly diagnosed and refractory MM patient samples. As shown in ***Fig. 1F*** (***see also Fig. S3***), the CC_50_ value of TCH-165 in cells from a newly diagnosed MM patient was 1.0μM (95% CI 0.60-1.51μM). Even the primary cells from a refractory MM patient were sensitive to TCH-165 at single digit micromolar potency (CC_50_ 8.1μM; 95% CI 7.08-9.03μM).

Consistent with this patient being refractory to bortezomib, bortezomib was completely ineffective against the isolated primary cells from the refractory MM patient (CC_50_ >50μM). Additional samples from larger populations will need to be investigated to determine whether 20S proteasome enhancers are generally effective in refractory patients. Experiments with both wild-type (WT) and bortezomib acquired (BTZ)-resistant human acute monocytic leukemia, THP-1 cells (kind gift from Dr. J. Cloos)^33^ exposed to TCH-165 (72h) at 5μM and 10μM (n=3) illustrated that the 20S proteasome enhancer readily overcomes acquired BTZ-resistance in cell culture (***Fig. 1G***).

### Gene expression profiling indicates limited changes in gene expression following TCH-165 treatment

The consequences of small molecule 20S activation in RPMI-8226 and CCRF-CEM cells on the expression of genes in key cancer pathways were evaluated using the NanoString nCounter system with the PanCancer Pathways™ panel ***(Fig. 2A-C)*** and analyzed using NanoString nSolver program and Ingenuity Pathway Analysis (IPA) (Qiagen). Even though this analysis method allows for accurate, reproducible comparisons of single experiments,^34^ this experiment was carried out twice in the RPMI-8226 cells (***Fig. 2C*** *see also* ***Fig. S4A***), which verified excellent reproducibility and accuracy of this assay. This multiplexed transcript detection assay analyzed the expression of 770 genes from 13 cancer-driving pathways. The gene profile of the vehicle control (DMSO) was compared to specific changes in gene expression induced by TCH-165 in CCRF-CEM and RPMI-8226 cells (***Fig. 2A-C***). Gene expression was normalized using the most stably expressed housekeeping genes in the panel in order to control for sample input variability and identified by the geNorm algorithm found within the nSolver advanced analysis module.^35^ The 22 housekeeping genes included for normalization are presented in Supplementary Figure S5.^35^ The heatmap generated by the correlation of pairwise analysis of gene expression reveals the intrinsic differences in gene expression profiles of the two cell lines, while showing the excellent agreement between repeated analysis carried out in the RPMI-8226 cell line, and the similarity between the vehicle controls in each cell line and at each time point ***(Fig. 2C*** *or by scatterplot in supplemental* ***Fig. S4B)***. Time and dose responsive changes at both 4h and 8h time points can also be observed, especially in the CCRF-CEM cell line.

Remarkably, the 20S proteasome enhancer affected only a few genes, with many of the differentially expressed genes being shared between the two cell lines in a concentration response manner. More genes were affected in the CCRF-CEM cell line than in RPMI-8226. Inspection of MYC expression in the two cell lines (***Fig. 3***) revealed significant differences between the two, with the RPMI-8226 cells having a much higher MYC expression, which remained largely unchanged in the presence of TCH-165, compared to CCRF-CEM cells, which showed a dose and time dependent decrease in MYC expression. Many genes down-regulated in both cell lines were designated MYC targets. All named genes in Figure 2A and Figure 2B indicate MYC targets identified through IPA and include TCF3, a key regulator of cell fate and development, especially in B-cell differentiation,^36^ and NOTCH1, also a key regulator of differentiation and proliferation, which is upregulated in many malignancies.^29^ Genes upregulated in both cells lines and all treatment points included DDIT3, the DNA damage inducible transcript 3, which enhances gene transcription via interaction with JUN and FOS and induces apoptosis following endoplasmic reticulum stress.^37^ Similarly, DDIT4, another DNA damage inducible transcript, which inhibits the mammalian target of rapamycin complex (mTROC1) was also consistently overexpressed. Of note, in the RPMI-8226 cells, the mRNA levels of the heat-shock protein, HSPA6, was also significantly upregulated by 51 and 55 fold, at 5 and 10 μM TCH-165, respectively. Interestingly, in the CCRF-CEM cells, where baseline HSPA6 mRNA levels were similar to RPMI8266 cells, no significant change was noted upon treatment with TCH-165.

### Pharmacokinetic properties and anti-tumor efficacy of 20S enhancer in *in vivo* xenograft

Following the cellular data, we initiated pharmacokinetic (PK) studies to determine plasma concentrations of TCH-165 following oral administration. CD-1 mice were exposed to 100 mg/Kg b.i.d. (oral gavage). The AUC was calculated using the trapezoidal method with mean values from data collected from 0-16 hours post-initial dose. Following an oral administration of TCH-165, the Cmax following the 1st dose was 932.3 ± 184.5 nM; with a Tmax at 2h. The Cmax following the 2nd dose was 1,434.0 ± 625.2 nM with a Tmax of 12h. The mean AUC (0-16h) was 13,975 h*nM/mL (***Fig. 4A and Fig. S6***). It is important to note that the ester moiety was (by LC-MS) not metabolically labile, likely due to its very sterically demanding location.^21^

**Figure 4:**
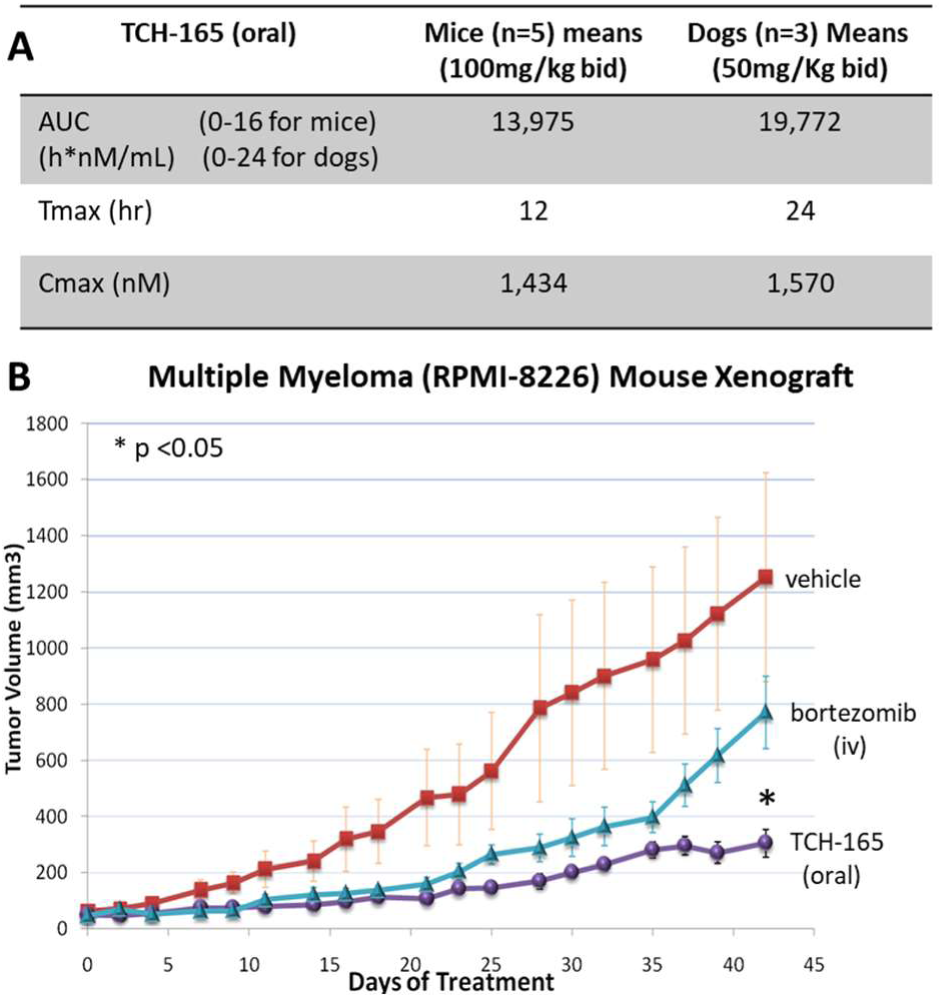
*In vivo* efficacy and tolerance of the proteasome activator, TCH-165. **(A)** Comparative analysis AUC, Tmax and Cmax of TCH-165 bid in mice (n=5) and dogs (n=3). **(B)** RPMI-8226 xenograft model in SCID mice treated with vehicle (3:7 (v/v) propylene glycol/5% dextrose), bortezomib (0.375 mg/Kg, iv, 3x/week), and TCH-165 (100mg/Kg, bid).

Considering that the concentration at which TCH-165 doubles the proteolytic activity of the 20S proteasome (***Fig. 1A***, AC_200_ = 1.5 μM) is similar to the effective concentrations against most cancer cell lines (***Fig. 1D***, CC_50_ 1-3 μM), as well as the maximum plasma concentration reached in mice (**Fig. 4A**), we evaluated the efficacy of this oral dose in an *in vivo* RPMI-8226 xenograft model. ***Figure 4B*** clearly shows that a dose of 100 mg/kg TCH-165, blocks tumor growth and is associated with minimal loss of body weight (<10%) indicating good tolerance (***Fig. S7***). Bortezomib (intravenous adminstration) was used as a positive control. Afer 42 days, the vehicle control group had an average tumor volume of 1253.4 (± 371.86) mm^3^, Bortezomib control 771.41 (±129.35) mm^3^ and TCH-165 304.43 (±49.50) mm^3^. Compared to the vehicle control, TCH-165 treatment by oral gavage resulted in a statistically significant reduction of tumor growth (75.71%, p<0.05) (***Fig. S8***).

### *In vivo* tolerance and target engagment of the 20S proteasome enhancer, TCH-165

Following these initial results of efficacy and tolerance, we examined TCH-165 for PK and tolerance in dogs. Adult male beagles received TCH-165 (oral capsule, 50mg/Kg, bid) for five consecutive days to match the AUC and Cmax as the (minimum) effective dose in mice (***Fig***.***4A-B***). This dose (50mg/Kg bid) resulted in an AUC (0-24h) of 19772 h*nM/mL with a Cmax of 1.6μM and a Tmax at 24h (***Fig. 4A, Fig. S9***). Clinical observations, body weight monitoring (<1% change), as well as standard panel of complete blood counts (***Fig. S10***) and clinical chemistry panel (***Fig. S11***) following five days of treatment identified no significant changes compared to pre-dose status. Target engagment studies using adult male beagles confirmed enhancement of proteasome activity following TCH-165 treatment. Peripheral blood mononuclear cells (PBMCs) were isolated from blood samples of untreated and treated (TCH-165, 1,920 mg BID, n=2 per group) canines. PBMCs were lysed and evaluated for hydrolysis of the CT-L substrate, Suc-LLVY-AMC and drug concentration using mass spectrometry, showing a 488% (1.4 μM) and 233% (1.1μM) signficantly increase substrate proteolysis (p<0.001), in the treated canines compared to untreated canines (***Fig. S12***). These studies indicate that the 20S proteasome enhancer, TCH-165, appears to be well tolerated *in vivo*, at concentrations where it significantly enhanced proteasome activity.

## DISCUSSION

Over-expression of c-MYC is one of the main oncogenic drivers in more than 50% of all human cancers, including most hematological malignancies. Hematologic malignancies are comprised of multiple subtypes including, multiple myeloma (MM), mantle cell lymphoma (MCL), acute myeloid leukemia (AML), histiocytic neoplasms, non-Hodgkin lymphoma, and many others.^38^ Despite the FDA approval of the proteasome inhibitors bortezomib,^39^ carfilzomib^40^ (and ixazomib for some patients)^41^ as front-line therapies for MM and MCL, patients still have poor prognosis, with a 5-year survival of 30-50%.^42^ Herein, we describe a novel mechanism to inhibit c-MYC activity through the stimulation of c-MYC degradation that may be therapeutically beneficial for not only hematological malignancies, but for all cancers that have enhanced c-MYC activity.

c-MYC is a transcription factor that activates the expression of many pro-proliferative genes following its dimerization to the myc-associated factor X (MAX).

c-MYC is a highly unstable protein that is regulated by the ubiquitin-proteasome system (UPS), following series of post-translational modification.^15,43^ Any dysregulation in c-MYC expression or in the posttranslation modification that delays or prevent its proteasome mediated degradation may result in upregulation of MYC regulated pro-proliferative genes. Importantly, c-MYC over-expression not only drives cancer growth, it also induces oncogenic transformation, chemoresistance and disease relapse.^44^

Considering the structurally disordered nature of c-MYC, we evaluated the efficacy of our previously reported 20S proteasome enhancer, TCH-165, on the induction of c-MYC degradation. We found that c-MYC protein levels were rapidly reduced (within 4 hours) in a concentration-dependent manner in the hematological cancer cell lines, RMPI-8226 and CCRF-CEM to TCH-165. Co-treatment with the proteasome inhibitor, bortezomib, prevented the TCH-165 induced degradation of c-MYC, thereby implicating the proteasome as the protease responsible. Furthermore, MYC mRNA remained unchanged upon TCH-165 treatment in RPMI-8226 cells, whereas a modest but dose dependent decrease was observed in CCRF-CEM cells. This modest difference in the c-MYC mRNA levels between the two cell lines may not be surprising considering the extraordinarily high level of c-MYC RNA in RPMI-8226 cells compared to the CCRF-CEM cells (**Fig. 3**), as well as the intrinsic differences between the cancer driving pathways of those cancer cell lines (**Fig. 2C** and **Fig. S3B**). Even though relatively few genes were affected by TCH-165 treatment, both cell lines differentially expressed multiple c-MYC target genes, including TCF3, NOTCH1, DDIT3, DDIT4, and VEGFA. These studies indicate that enhanced 20S proteasome activity primarily induces the degradation of excessive (for example, overexpressed) IDPs and that “normal” levels of IDPs may not be significantly affected, thereby initially only affecting a relatively small set of proteins, and consequently gene targets.

Proteasome inhibitors including bortezomib are the initial treatment option in various hematologic malignancies, unfortunately many patients relapse or may not respond to treatment at all (refractory patients). Overcoming resistance is critical in the search for new therapeutic approaches. The 20S proteasome enhancer, TCH-165, was found to be effective in bortezomib resistant cell lines, as well as in primary cells from a patient who did not respond (refractory) to bortezomib treatment. Whether or not the decrease in c-MYC protein by 20S proteasome enhancement is responsible for overcoming this resistance is not currently known, but an intriguing possibility.

The translational efficacy of the 20S proteasome enhancer was evaluated in an RPMI-8226, multiple myeloma xenograft model. TCH-165 when administered to mice at 100 mg/kg twice daily resulted in plasma concentrations approximating the AC_200_. At this dose, a significant inhibition of tumor growth was observed (***Fig. 4A-B***) in the mice multiple myeloma xenograft model. The treatment was well tolerated by the mice, with <10% loss of body weight, resulting in 76% reduction of tumor growth. More detailed tolerance of 20S enhancement by TCH-165 was examined by approximating the AUC and Cmax of the mouse study in beagles (n = 3) with TCH-165 treatment for five consecutive days (oral capsule, 50mg/Kg, bid). Clinical observation, body weight (<1% change), as well as the standard complete blood count (37 criteria) and clinical chemistry (29 criteria) following this five-day tolerance study identified no significant changes compared to pre-treatment evaluations. These studies demonstrated the first translational anti-tumor efficacy and acceptable tolerance *in vivo* of the 20S enhancer, TCH-165, and support further preclinical investigations in MM disease models that more accurately recapitulate hematological diseases.^45^ Overall, the active concentration at which TCH-165 doubles 20S proteasome activity (AC_200_ 1.5μM) corresponded well to the effective concentration required to induce 50% cell death (CC_50_) in various cell culture, and corresponded to the Cmax obtained *in vivo*, and required to inhibit tumor growth.

Our data combined indicate that enhancing the 20S proteolytic activity, using TCH-165, targets excessive levels of disease-driving disordered proteins, with relatively few changes to the cellular proteome. 20S proteasome enhancement reduces c-MYC protein levels, which may be responsible for its anti-cancer efficacy, however other disordered protein targets cannot be excluded. Gene expression of various cancer pathways showed a decrease in expression of several tumor promoting genes and an increase in pro-apoptotic genes in cancer cell lines. Importantly, the 20S proteasome enhancer was well tolerated both *in vitro* and *in vivo*, thereby validating this new approach for further therapeutic assessment for various IDP-driven proteotoxic disorders.

## EXPERIMENTAL PROCEDURES

### *In vitro* 20S proteasome mediated degradation of c-MYC

Varying concentrations of purified 20S proteasome (final concentrations ranging from 0-240 nM) in assay buffer (50 mM Tris-HCl buffer, pH 7.8) were incubated with purified c-MYC (final concentration 0.5 µM) for 48 hours at 37 °C. Following incubation, GAPDH was added to a final concentration of 0.5 µM as a loading control, and the samples were boiled at 90 °C with 5X SDS loading buffer. The samples were resolved on a 4-20% Tris/glycine gel and transferred to a PVDF membrane. The membrane was probed with anti c-MYC (Cell signaling; cat#5605) and anti-GAPDH-HRP (Santa Cruz; sc-47724) antibodies.

### Cell culture

RPMI-8226 and CCRF-CEM cells from ATCC were maintained in RPMI medium supplemented with 10% Fetal Bovine Serum, and 100U/mL Penicillin/Streptomycin, at 37°C with 5% CO_2_. DMSO was used as compound vehicle or vehicle control at 1% final concentration for both cell culture and enzyme assay.

### Primary cell culture and cell viability

Bone marrow aspirates were obtained by Dr. Isaac Daniel or by Dr. Omar Alkharabsheh, following standard protocol. WBCs were separated from RBCs by ficoll density gradient. Multiple myeloma cells were then isolated from WBCs using human CD138 affinity magnetic beads (Miltenyi Biotec; cat#130-093-062) following the manufacturer’s protocol. Cells were cryopreserved in 10% DMSO in FBS. Revived cells were maintained in RPMI medium with 10% FBS for 10 days prior to treatment. Primary or established cells (10,000/well) were seeded in 96-well plate in 100μL medium and treated with TCH-165 or Bortezomib for 72h. Cells were equilibrated to RT and CellTiter-Glo (Promega) solution (100μL) added and incubated with shaking for 10 minutes at RT. Assay plate was then allowed to equilibrate for 5 minutes at RT and luminescent readings taken on a SpectraMax M5^e^. Data are presented as a percentage of the vehicle control for each experimental condition, after background subtraction. Error bars represent SD for triplicate wells.

### Proteasome activity in purified protein assay

Proteasome activity were carried out as previously reported.^25^

### c-MYC degradation in RPMI-8226 and CCRF-CEM cells

CCRF-CEM or RPMI-8226 cells were grown to approximately 80% confluency in a T-75 flask. Cells were treated with either vehicle (DMSO) or TCH-165 at the indicated concentrations, for 4h. In experiments involving bortezomibs, cells were pretreated with bortezomib (5μM) for 1h before adding TCH-165 or vehicle for a further 4h. Cells were pelleted and washed with chilled PBS buffer (2X) and resuspended in chilled RIPA buffer supplemented with sigmafast protease inhibitor cocktail (Sigma Aldrich). Total protein was quantified by bicinchoninic acid assay (BCA assay, Thermofisher Scientific), normalized to 2mg/mL and boiled with 5X SDS loading buffer. Equal amounts of lysates (30μg) were resolved on a 4-20% Tris/glycine gel and transferred to a PVDF membrane. Membrane was probed with anti c-MYC (Cell signaling; cat#5605) and anti-GAPDH-HRP (Santa Cruz; sc-47724 HRP) anrtibodies.

### Gene expression profiling of CCRF-CEM and RPMI-8226 cells

CCRF-CEM cells or RPMI-8226 cells were treated with vehicle, TCH-165 (5μM or 10μM) for 4h or 8h. Total RNA was obtained using *mir*Vana™ miRNA Isolation Kit, with phenol (Thermofisher Sientific; AM1560). Total mRNAs were quantified by Qubit (Thermofisher Scientific; Q32855) and gene expression evaluated against the PanCancer Pathways panel using standard NanoString protocol.

### Pharmacokinetic assessment of TCH-165

PK study was carried out with CD-1 mice (males, 6-8 weeks of age, n=3 per treatment group) weighing between 24.9 g to 33.7 g immediately prior to dosing. TCH-165 (100 mg/kg) was administered by oral gavage, as a homogeneous suspension in a 3:7 (v/v) propylene glycol: 5% dextrose as vehicle. A second dose of 100 mg/kg TCH-165 was administered approximately 8 hours post-initial gavage. Blood was collected at 0.5h, 1h, 2h, 4h, 8h, 9h, 10h, 12h, and 16h post-initial gavage in Li-Heparin tubes. Plasma was separated by centrifugation and analyzed using mass spectrometry.

### Pharmacokinetic assessment and tolerance in dogs

Adult beagle dogs (male, 12 months of age, n=3) weighing approximately 10Kg received a single oral dose of test compound at 500 mg bid delivered in capsule form. Blood (approximately 2.5 mL) was collected into lithium heparin tubes at 0, 1h, 2h, 4h, 8h, and 24h post-day 1 and post-day 5 dose for PK assessment using mass spectrometry. Blood (approximately 2 mL) was also collected and divided into EDTA and standard clot tubes at day 6 post-dose for determination of CBC and Clinical chemistry parameters, respectively. Clinical observations were conducted daily on non-dosing days and twice on dosing days.

### Proteasome Activity in Canine PBMC Lysate

Canine blood samples obtained in BD Vacutainer® CPT™ mononuclear cell preparation tubes (treated with sodium citrate) and were used to isolate PBMCs. For PBMC isolation, sample tubes were inverted five times and equilibrated for the centrifuge with sterile PBS buffer. Tubes were centrifuged at 1800 xg with acceleration set at 5 and break set to 0 at room temperature for 30 minutes. Half of the plasma layer was discarded. Using a sterile Pasteur pipette the buffy coat was collected and put into a 15 mL conical tube and sterile PBS buffer was added to 10 mL. The conical tubes were centrifuged at 300 xg for 10 minutes and the supernatant was removed. Cells were resuspended in 150 µL of lysis buffer (50 mM Tris-HCl, 2 mM Na_2_ATP, 5 mM MgCl_2_, 0.5 mM EDTA, 10% glycerol) and lysed by vortex. Samples were centrifuged for 20 minutes at 14,000 xg. Total protein concentration of the supernatant was determined using bicinchoninic acid assay (BCA assay), and the samples were normalized to 1 mg/mL. Samples were diluted to 0.036 µg/µL in assay buffer (38 mM Tris, 100 mM NaCl, pH 7.8) and 140 µL of the diluted sample was added to three wells of a black, clear-bottom 96-well plate. Substrate stock solution (10 µL of 375 µM Suc-LLVY-AMC in assay buffer) was added to each well. Fluorescence was measured by kinetic readings taken every 5 minutes at 37 °C, at 380/440 nm for 1 h.

### RPMI-8226 xenograft tumor model

CB17-SCID mice (Female, 3-4 weeks of age) were injected subcutaneously with 1.2 ⨯ 10^7^ cells per animal. Treatment was initiated when tumors reached ∼50mm^3^. Tumor dimensions were measured three times weekly using digital calipers. Body weights were measured three times weekly. TCH-165 (150mg/kg on day 1, reduced to 100mg/Kg on day 2 and for the remainder of the study), was prepared as a homogeneous suspension in a 3:7 (v/v) propylene glycol: 5% dextrose vehicle. Bortezomib (0.375 mg/Kg-day 0, 0.18 mg/Kg-day 2, 0.09 mg/Kg day 5 and beyond) was given 3X per week intravenously. Bortezomib was reduced to 0.09 mg/kg per treatment for the remainder of the study, due to weight loss at higher doses.

## SUPPLEMENTAL INFORMATION

The supporting Information includes Supplemental Experimental Procedures and twelve figures referenced in the article.

## ACKNOWLEDGMENTS

The authors thank E. Crisovan and C. Harris from the Genomics Core Facilities, Teresa Krieger-Burke from the *In Vivo* facility, and Bilal Aleiwi from the Medicinal Chemistry Core Facility for their assistance and expert analysis. Financial support for this work was provided by the National Institute of Allergy and Infectious Diseases (1R21AI117018-01A1) and the National Institute of General Medical Sciences (T32GM092715) of the National Institutes of Health. The authors also gratefully acknowledge financial support from the international myeloma foundation (IMF) Brian D. Novis Grant and Michigan State University (SPG and CTSI grant programs), the Michigan Translational Research and Commercialization innovation hub and from the Robert and Jean P. Schultz Biomedical Research Endowment Award, NIEHS Toxicology Training Grant T32 ES007255.

## AUTHOR CONTRIBUTIONS

The manuscript was written through contributions of all authors.

