## supplemental file for "Enhancing c-MYC degradation via 20S proteasome activation induces *in vivo* anti-tumor efficacy"

**Fig. S12** Target engagement study in treated and untreated dogs……………………………………………..14

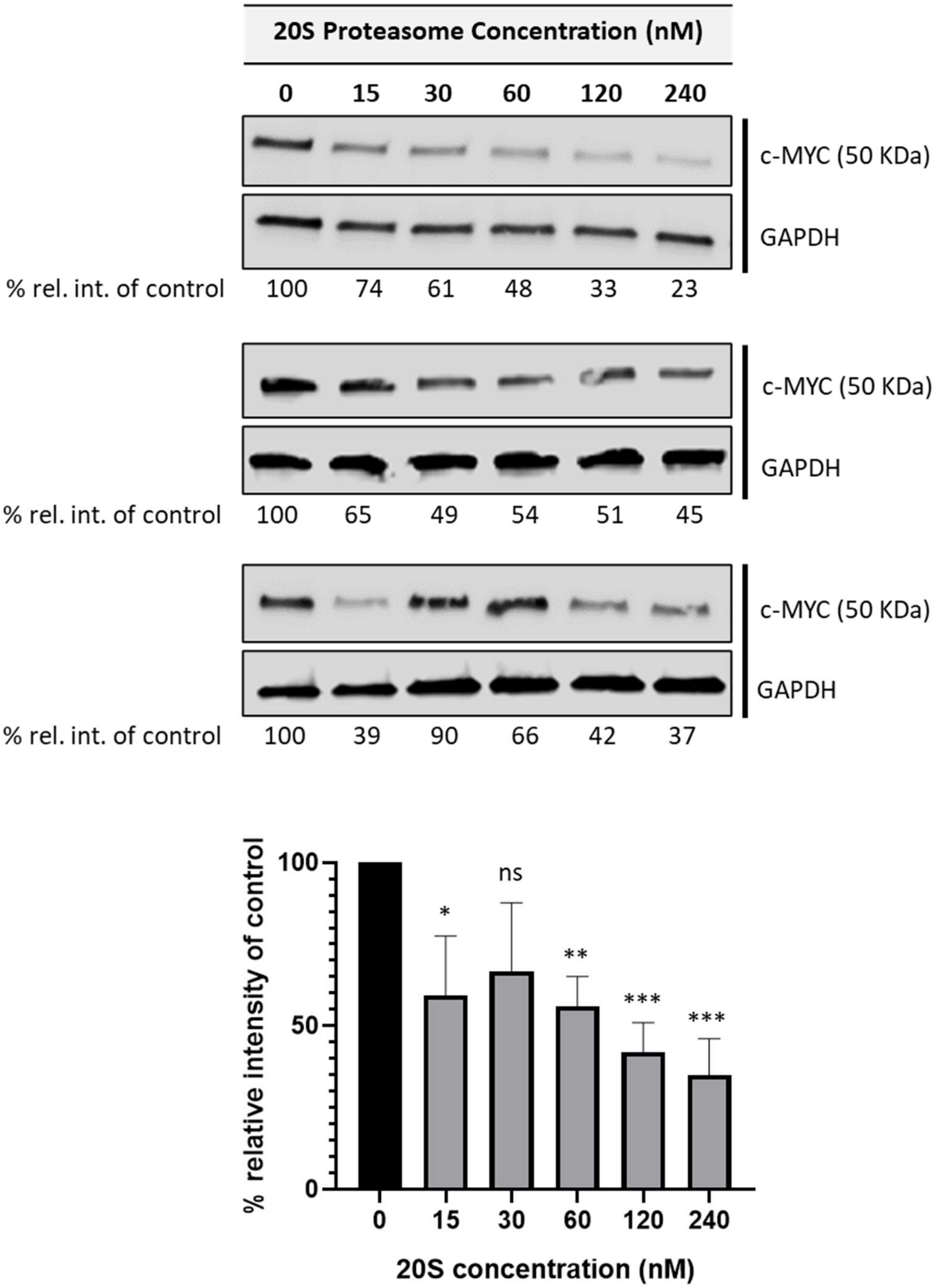

**Fig. S1: c-MYC degradation by the 20S proteasome (n=3)**: Immunoblot of c-MYC exposed to increasing concentrations of purified 20S proteasome after 48hr exposure.

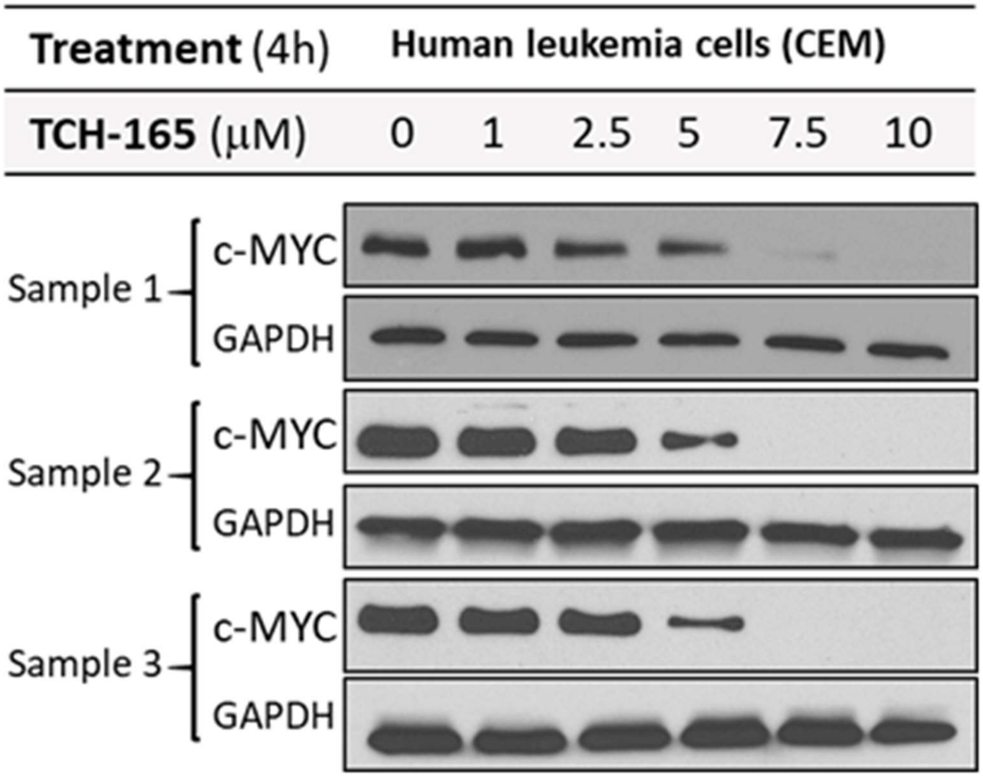

**Fig. S2: c-MYC degradation immunoblots (n=3)**: Three different immunoblot of CCRF-CEM cell lysates, showing reproducible, concentration dependent reduction of c-MYC in human leukemia cells upon 4h treatment with various concentrations of TCH-165.

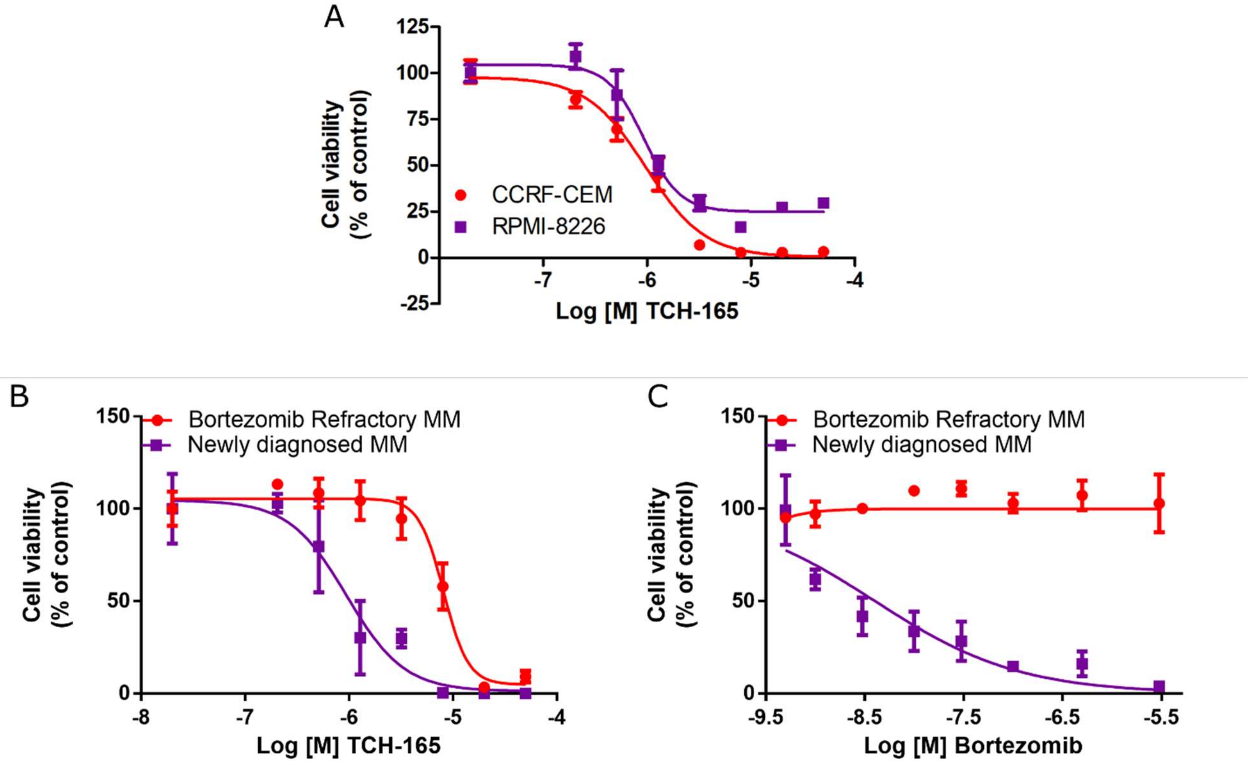

**Fig. S3: Concentration response curves for Fig. 1F**. **(A)** Viability of lymphoblastic leukemia (CCRF-CEM) and multiple myeloma (RPMI-8226) cells following treatment with TCH-165 for 72h. Multiple myeloma cells isolated from bone marrow aspirates of a newly diagnosed patient or a patient who is inherently resistant to bortezomib treatment were treated with TCH-165 **(B)** or Bortezomib **(C)** and cell viability measured after 72h

**A**

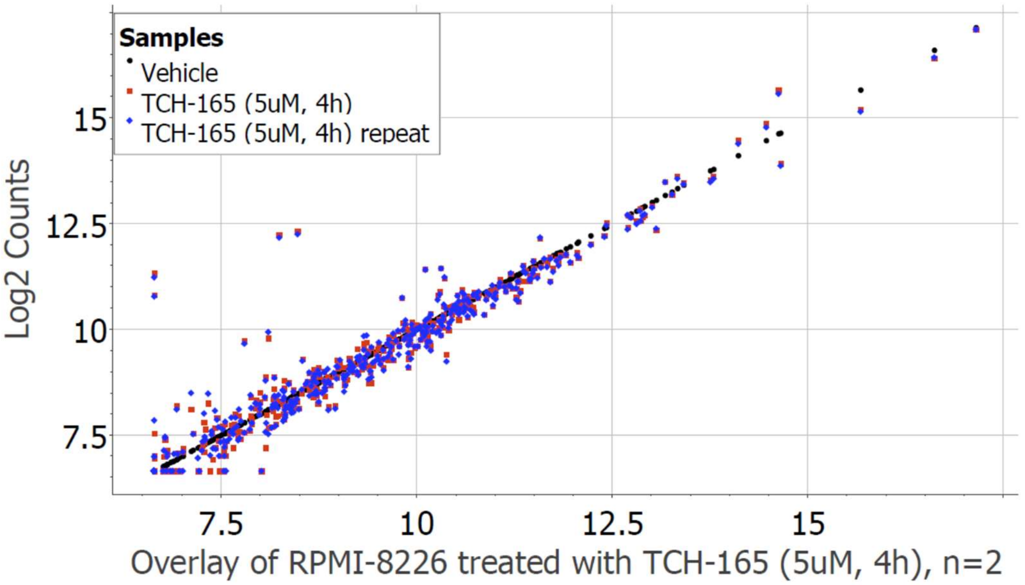

**B**

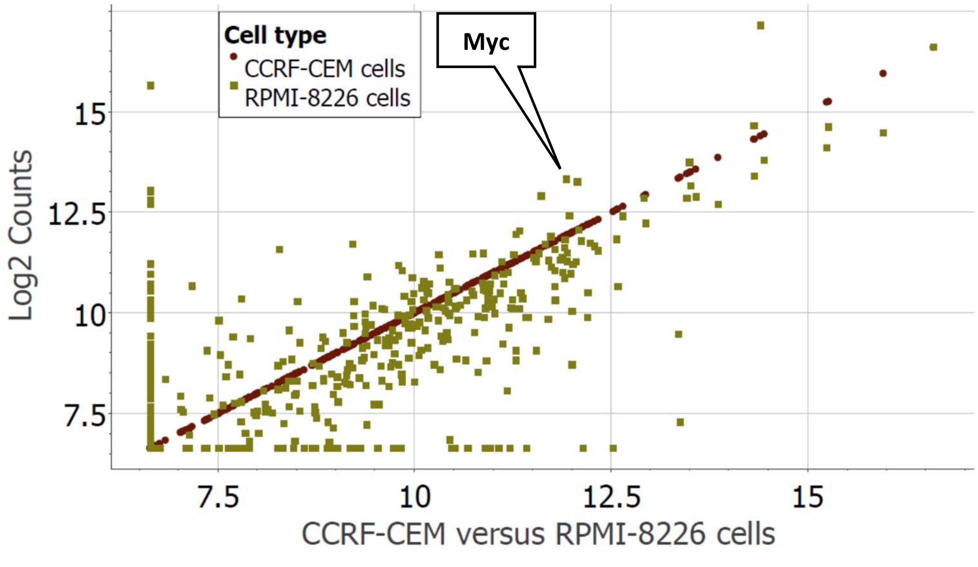

**Fig. S4**: **Gene expression profiles. A**. Scatter plot of the gene expression profile of CCRF-CEM cells compared to RPMI-8226 cells treated with vehicle for 4 hours. **B**. Scatter plot of RPMI-8226 cells treated with vehicle (4h) and treated withTCH-165 (5 μM, 4h), in two independent experiments (n=2).

| Gene Name | Order selected by geNorm | SD after normalization |
|---|---|---|
| SLC4A1AP-mRNA | 1 | 0.133 |
| DHX16-mRNA | 2 | 0.106 |
| CNOT4-mRNA | 3 | 0.12 |
| PRPF38A-mRNA | 4 | 0.0707 |
| VPS33B-mRNA | 5 | 0.0907 |
| EIF2B4-mRNA | 6 | 0.113 |
| COG7-mRNA | 7 | 0.102 |
| MRPS5-mRNA | 8 | 0.141 |
| ACAD9-mRNA | 9 | 0.111 |
| SF3A3-mRNA | 10 | 0.188 |
| PIK3R4-mRNA | 11 | 0.18 |
| RBM45-mRNA | 12 | 0.214 |
| SAP130-mRNA | 13 | 0.187 |
| ZNF384-mRNA | 14 | 0.205 |
| MTMR14-mRNA | 15 | 0.16 |
| FTSJ2-mRNA | 16 | 0.181 |
| PIAS1-mRNA | 17 | 0.189 |
| NOL7-mRNA | 18 | 0.2 |
| TMUB2-mRNA | 19 | 0.198 |
| CNOT10-mRNA | 20 | 0.21 |
| GPATCH3-mRNA | 21 | 0.223 |
| TRIM39-mRNA | 22 | 0.237 |
| DNAJC14-mRNA | discarded | 0.332 |
| ZKSCAN5-mRNA | discarded | 0.345 |
| EDC3-mRNA | discarded | 0.353 |
| ERCC3-mRNA | discarded | 0.367 |
| FCF1-mRNA | discarded | 0.375 |
| NUBP1-mRNA | discarded | 0.375 |
| TLK2-mRNA | discarded | 0.391 |
| AGK-mRNA | discarded | 0.411 |
| TTC31-mRNA | discarded | 0.426 |
| ZNF143-mRNA | discarded | 0.456 |
| C10orf76-mRNA | discarded | 0.518 |
| AMMECR1L-mRNA | discarded | 0.518 |
| ZC3H14-mRNA | discarded | 0.585 |
| CC2D1B-mRNA | discarded | 0.611 |
| USP39-mRNA | discarded | 0.631 |
| HDAC3-mRNA | discarded | 0.678 |
| ZNF346-mRNA | discarded | 0.763 |
| DDX50-mRNA | discarded | 1.13 |

**Fig. S5**. House keeping genes used for normalization of gene expression

| Group | PK Timepoint (h) Post-Initial Gavage | Animal # | Plasma Concentration (nM) | Mean Plasma Concentration (nM) | ± Standard Deviation |
|---|---|---|---|---|---|
| 1 | 0.5 | 1 | 447.6 | 355.1 | 181.5 |
| | | 2 | 146.0 | | |
| | | 3 | 471.6 | | |
| 2 | 1 | 4 | 198.0 | 620.8 | 398.6 |
| | | 5 | 989.6 | | |
| | | 6 | 674.8 | | |
| 3 | 2 | 7 | 819.2 | 932.3 | 184.5 |
| | | 8 | 832.4 | | |
| | | 9 | 1145.2 | | |
| 4 | 4 | 10 | 764.0 | 678.9 | 323.3 |
| | | 11 | 951.2 | | |
| | | 12 | 321.6 | | |
| 5 | 8 | 22 | 885.6 | 845.9 | 373.6 |
| | | 23 | 1198.0 | | |
| | | 24 | 454.0 | | |
| 6 | 9 | 16 | 1321.2 | 1170.9 | 130.6 |
| | | 17 | 1107.2 | | |
| | | 18 | 1084.4 | | |
| 7 | 10 | 19 | 646.0 | 911.6 | 282.7 |
| | | 20 | 880.0 | | |
| | | 21 | 1208.8 | | |
| 8 | 12 | 13 | 2138.4 | 1434.0 | 625.2 |
| | | 14 | 1218.8 | | |
| | | 15 | 944.8 | | |
| 9 | 16 | 25 | 840.4 | 471.1 | 330.5 |
| | | 26 | 203.2 | | |
| | | 27 | 369.6 | | |

**Fig. S6:** Pharmacokinetic data in mice. TCH-165 plasma concentrations after oral gavage 3:7 (v/v) propylene glycol: 5% D5W vehicle of male CD-1 mice.

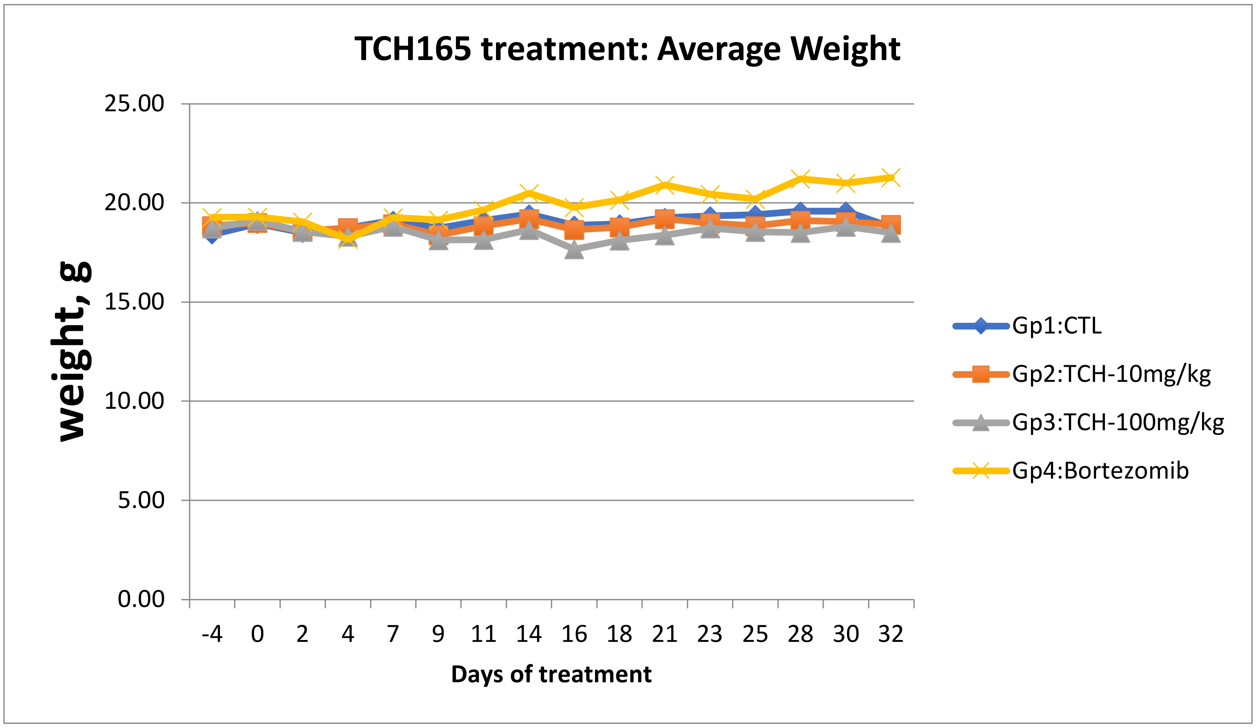

| Days: | -4 | 0 | 2 | 4 | 7 | 9 | 11 | 14 | 16 | 18 | 21 | 23 | 25 | 28 | 30 | 32 |
|---|---|---|---|---|---|---|---|---|---|---|---|---|---|---|---|---|
| Gp1:CTL | 18.42 | 18.95 | 18.50 | 18.73 | 19.12 | 18.73 | 19.11 | 19.43 | 18.86 | 18.93 | 19.26 | 19.34 | 19.41 | 19.58 | 19.57 | 18.77 |
| Gp2:TCH-10mg/kg | 18.8 | 19.0 | 18.6 | 18.7 | 18.9 | 18.4 | 18.8 | 19.2 | 18.6 | 18.8 | 19.2 | 19.0 | 18.8 | 19.1 | 19.0 | 18.9 |
| Gp3:TCH-100mg/kg | 18.72 | 19.11 | 18.54 | 18.31 | 18.80 | 18.13 | 18.13 | 18.64 | 17.67 | 18.10 | 18.37 | 18.72 | 18.55 | 18.50 | 18.80 | 18.49 |
| Gp4:Bortezomib | 19.28 | 19.29 | 19.04 | 18.17 | 19.25 | 19.14 | 19.65 | 20.49 | 19.77 | 20.14 | 20.91 | 20.44 | 20.20 | 21.20 | 21.00 | 21.29 |

**Fig. S7**: Weights of mice during tumor study.

| Mean | Days | -17 | -14 | -11 | -7 | -3 | 0 | 2 | 4 | 7 | 9 | 11 | 14 | 16 | 18 | 21 | 23 | 25 | 28 | 30 | 32 | 35 | 37 | 39 | 42 |
|---|---|---|---|---|---|---|---|---|---|---|---|---|---|---|---|---|---|---|---|---|---|---|---|---|---|
| **Gp1: Control** | | 10.60 | 15.43 | 43.95 | 41.76 | 38.49 | 63.25 | 70.62 | 90.73 | 137.86 | 163.48 | 212.01 | 241.02 | 319.30 | 346.02 | 466.65 | 478.21 | 561.67 | 786.61 | 841.01 | 900.52 | 958.86 | 1026.87 | 1122.13 | 1253.46 |
| **Gp2:TCH-10mg/kg** | | 9.68 | 16.32 | 30.88 | 45.85 | 36.36 | 58.33 | 55.23 | 70.72 | 89.68 | 98.72 | 103.98 | 138.48 | 148.48 | 230.71 | 281.51 | 343.57 | 395.40 | 569.20 | 739.04 | 733.71 | 867.08 | 1028.36 | 1046.77 | 1395.61 |
| **Gp3:TCH-100mg/kg** | | 11.18 | 19.37 | 32.31 | 37.50 | 38.99 | 49.70 | 47.20 | 54.74 | 73.95 | 74.03 | 79.59 | 84.60 | 96.08 | 112.96 | 107.12 | 142.69 | 145.59 | 169.57 | 177.03 | 198.99 | 239.73 | 252.08 | 270.05 | 304.43 |
| **Gp4:Bortezomib** | | 10.30 | 8.87 | 18.10 | 28.61 | 37.38 | 47.25 | 69.28 | 52.54 | 62.66 | 64.45 | 104.43 | 122.46 | 126.51 | 137.34 | 158.57 | 204.03 | 264.35 | 287.39 | 323.49 | 363.80 | 396.01 | 510.98 | 617.56 | 771.41 |

**Fig. S8**: Tumor volume data from TCH-165 treated RPMI-8226 xenograft model using SCID mice.

**A.** Plasma concentration of TCH-165 at Day 1 and Day 5.

| | Dog 1 | Dog 2 | Dog 3 | Mean | SD | N |
|---|---|---|---|---|---|---|
| **Pre-dose** | 0 | 0 | 0 | 0 | 0 | 3 |
| **1hr SD1** | 25 | 0 | 23 | 16 | 14 | 3 |
| **2hr SD1** | 103 | 63 | 160 | 109 | 49 | 3 |
| **4hr SD1** | 183 | 121 | 97 | 134 | 44 | 3 |
| **8hr SD1** | 130 | 41 | 45 | 72 | 50 | 3 |
| **9hr SD1** | 534 | 453 | 33 | 340 | 269 | 3 |
| **10hr SD1** | 1444 | 1390 | 239 | 1024 | 681 | 3 |
| **24hr SD1** | 909 | 2199 | 1603 | 1570 | 645 | 3 |
| **1hr SD5** | 593 | 961 | 750 | 768 | 184 | 3 |
| **2hr SD5** | 605 | 881 | 792 | 759 | 141 | 3 |
| **4hr SD5** | 747 | 1017 | 595 | 786 | 214 | 3 |
| **8hr SD5** | 756 | 782 | 387 | 641 | 221 | 3 |
| **9hr SD5** | 841 | 873 | 350 | 688 | 293 | 3 |
| **10hr SD5** | 1622 | 1462 | 733 | 1272 | 474 | 3 |
| **24hr SD5** | 546 | 1225 | 1135 | 969 | 369 | 3 |

**B.** PK of TCH-165 at Day 1 and Day 5

| | **Dog 1** | **Dog 2** | **Dog 3** | Mean |
|---|---|---|---|---|
| $AUC_{(0-24)}$ Day 1 (nM) | 18781 | 2631 | 13713 | 19772 |
| Cmax Day 1 (nM) | 1444 | 2199 | 1603 | 1570 |
| Tmax Day 1 (nM) | 10 | 24 | 24 | 24 |
| $AUC_{(0-24)}$ Day 5 (nM) | 22163 | 27221 | 18108 | 22494 |
| Cmax Day 5 (nM) | 1622 | 1462 | 1135 | 1272 |

**Fig. S9**: Pharmacokinetic data in dogs: **A.** Plasma concentration (nM) of TCH-165 following oral gavage (500mg BID) at Day 1 (SD1) and Day 5 (SD5). **B.** Pharmacokinetic parameter of TCH-165 (500mg BID).

| | | Pre Dose | | | Day 6 | | |
|---|---|---|---|---|---|---|---|
| | | Dog 001 | Dog 002 | Dog 003 | Dog 001 | Dog 002 | Dog 003 |
| **Hemolysis** | | Normal | Normal | Normal | Normal | Normal | Normal |
| **Lipemia** | | Normal | Normal | Normal | Normal | Normal | Normal |
| **Icterus** | | Normal | Normal | Normal | Normal | Normal | Normal |
| **Total Protein** | *g/dL* | 6.7 | 6.4 | 6.8 | 7 | 6.4 | 6.4 |
| **RBC** | *x 10.e6/uL* | 7.1 | 6.8 | 7 | 7.3 | 7.4 | 6.2 |
| **Hgb** | *g/dL* | 16.1 | 15.9 | 16.3 | 16.8 | 17.6 | 14.9 |
| **Hct** | *%* | 47 | 46 | 47 | 50 | 50 | 44 |
| **HCT Spun** | *%* | 48 | 44 | 45 | 50 | 51 | 45 |
| **MCV** | *fL* | 66 | 67 | 68 | 68 | 69 | 70 |
| **MCH** | *pg* | 23 | 23 | 23 | 23 | 24 | 24 |
| **MCHC** | *g/dL* | 34.0 | 35.0 | 34.0 | 34.0 | 35.0 | 34.0 |
| **CHCM** | *g/dL* | 33* | 34 | 34 | 32 * | 33 | 32 * |
| **RDW** | *%* | 13** | 13** | 13** | 13 ** | 12 | 12 |
| **Platelet** | *x10.e3/uL* | 287 | 269 | 188 | 269 | 242 | 151* |
| **MPV** | *fL* | 10.4 | 9.4 | 13.7 | 11 | 10.4 | 15.1 ** |
| **WBC** | *x10.e3/uL* | 9.7a | 10.9 | 8.2 | 9.4 | 10.8 | 7.9 |
| **Seg Neut #** | *x10.e3/uL* | 5.7 | 7.1 | 5.7 | NA | NA | NA |
| **Neutrophil #** | *x10.e3/uL* | NA | NA | NA | 5.9 | 6.1 | 4.5 |
| **Band Neutrophil #** | *x10.e3/uL* | 0.1 | 0.2** | 0.1 | NA | NA | NA |
| **Lymphocyte #** | *x10.e3/uL* | 2.8 | 3.1 | 1.6 | 2.6 | 3.8** | 2.5 |
| **Monocyte #** | *x10.e3/uL* | 0.6 | 0.3 | 0.8 | 0.6 | 0.5 | 0.6 |
| **Eosinophil #** | *x10.e3/uL* | 0.5 | 0.2 | 0 | 0.3 | 0.3 | 0.2 |
| **Basophil #** | *x10.e3/uL* | 0 | 0 | 0 | NA | NA | NA |
| **LUC #** | *x10.e3/uL* | NA | NA | NA | NA | NA | 0.03 |
| **Neutrophil Pct** | *%* | NA | NA | NA | 62.9 | 56.7 | 56.8 |
| **Seg Neut Pct** | *%* | 59 | 65 | 70 | NA | NA | NA |
| **Band Neut Pct** | *%* | 1 | 2 | 1 | NA | NA | NA |
| **Lymphocyte Pct** | *%* | 29 | 28 | 19 | 27.1 | 35.2 | 31.3 |
| **Monocyte Pct** | % | 6 | 3 | 10 | 6.1 | 4.8 | 7.8 |
| **Eosinophil Pct** | % | 5 | 2 | 0 | 3 | 2.8 | 3.2 |
| **Basophil Pct** | % | 0 | 0 | 0 | NA | NA | NA |
| **LUC Pct** | % | 1 | NA | NA | 0.4 | 0.2 | 0.3 |
| **NRBC** | /100 WBC | 1 | NA | NA | NA | NA | NA |
| **NRBC #** | *x10.e3/uL* | 0.1 | NA | NA | NA | NA | NA |
| **Reactive Lymphs** | | NA | NA | NA | Present | Present | NA |
| **Platelet Clump** | | NA | NA | present | Present | Present | Present |
| **Platelet Comment** | | NA | NA | b | b | b | b |

***** **Low Result**

**** High Result**

**a - WBC corrected for nucleated RBCs**

**b - Platelet concentration should be considered a minimum value and the MPV may be falsely increased due to platelet clumping.**

**Fig. S10**: Blood count panel of treated versus untreated dogs

| | | Pre-Dose | | | 24h Post Day 5 AM Dose | | |
|---|---|---|---|---|---|---|---|
| | | Dog 001 | Dog 002 | Dog 003 | Dog 001 | Dog 002 | Dog 003 |
| **Urea Nitrogen** | *mg/dL* | 20 | 16 | 21 | 17 | 14 | 18 |
| **Creatinine** | *mg/dL* | 1 | 0.8 | 0.9 | 0.8 | 0.6 | 0.7 |
| **Sodium** | *mmol/L* | 148 | 148 | 147 | 146 | 144 | 146 |
| **Potassium** | *mmol/L* | 4.5 | 4.4 | 4.7 | 4.7 | 4.6 | 4.4 |
| **Chloride** | *mmol/L* | 111 | 111 | 106 | 110 | 112 | 110 |
| **TCO2** | *mmol/L* | 23 | 24 | 27 | 22 | 18 | 22 |
| **Anion Gap** | *mmol/L* | 18 | 17 | 19 | 19 | 19 | 18 |
| **Na/K Ratio** | | 33 | 34 | 31 | 31 | 31 | 33 |
| **Osmolarity Calc** | *mOs/L* | 308 | 307 | 306 | 303 | 298 | 303 |
| **Glucose** | *mg/dL* | 88 | 92 | 87 | 91 | 83 | 87 |
| **Calcium** | *mg/dL* | 10.1 | 10.0 | 10.2 | 10.3 | 10.3 | 9.9 |
| **Magnesium** | *mg/dL* | 1.9 | 1.9 | 2 | 1.8 | 1.6 * | 1.7 |
| **Phosphorus** | *mg/dL* | 4.4 | 4.6 | 4.9 | 4.3 | 4.4 | 4.3 |
| **Iron** | *ug/dL* | 126 | 147 | 108* | 276** | 284** | 307** |
| **Total Protein** | *g/dL* | 5.9 | 5.7 | 5.9 | 5.4 | 5.0* | 5.0* |
| **Albumin** | *g/dL* | 3.3 | 3.2 | 3.2 | 3.1 | 3 | 2.8 |
| **Globulin Calc** | *g/dL* | 2.6 | 2.5 | 2.7 | 2.3 | 2.0* | 2.2* |
| **Total Bili** | *mg/dL* | 0.2 | 0.2 | 0.2 | 0.1 | 0.2 | 0.1 |
| **Direct Bili** | *mg/dL* | 0 | 0 | 0 | 0 | 0 | 0 |
| **Indirect Bili** | *mg/dL* | 0.2 | 0.2 | 0.2 | 0.1 | 0.2 | 0.1 |
| **Amylase** | *U/L* | 612 | 457 | 590 | 495 | 505 | 516 |
| **ALP** | *U/L* | 59 | 32 | 35 | 66 | 71 | 47 |
| **ALT** | *U/L* | 19* | 34 | 24 | 25 | 44 | 23 |
| **AST** | *U/L* | 26 | 37 | 34 | 23 | 32 | 24 |
| **Chol** | *mg/dL* | 111 | 330** | 125 | 184 | 140 | 190 |
| **CK** | *U/L* | 180 | 136 | 203 | 78 | 241** | 83 |
| **Hemolysis** | | Normal | Normal | Normal | Normal | Slight | Normal |
| **Icterus** | | Normal | Normal | Normal | Normal | Normal | Normal |
| **Lipemia** | | Normal | Normal | Normal | Normal | Normal | Normal |

* **Low Result**

** **High Result**

**Fig. S11**: Clinical chemistry panel of treated versus untreated dogs

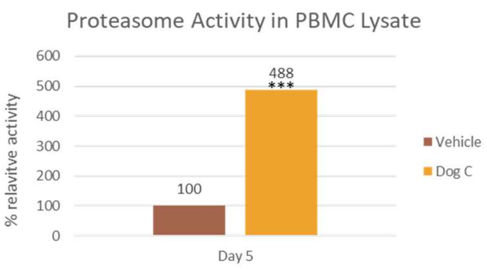

| Dog 1 | | | | | |
|---|---|---|---|---|---|
| Trial 1 | | Trial 2 | | Trial 3 | |
| Vehicle | Day 5 | Vehicle | Day 5 | Vehicle | Day 5 |
| 0.019 | 0.081 | 0.017 | 0.087 | 0.016 | 0.085 |
| 0.018 | 0.079 | 0.015 | 0.083 | 0.020 | 0.089 |
| 0.017 | 0.077 | 0.015 | 0.086 | 0.017 | 0.084 |
| **0.018** | **0.079** | **0.016** | **0.085** | **0.018** | **0.086** |
| 4.39 | | 5.45 | | 4.87 | |

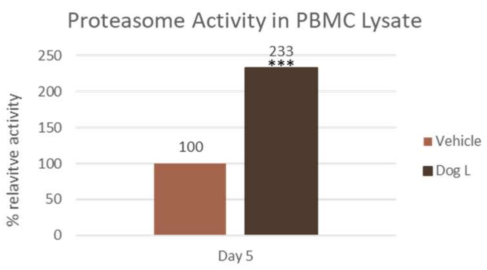

| Dog 2 | | | | | |
|---|---|---|---|---|---|
| Trial 1 | | Trial 2 | | Trial 3 | |
| Vehicle | Day 5 | Vehicle | Day 5 | Vehicle | Day 5 |
| 0.038 | 0.082 | 0.041 | 0.093 | 0.044 | 0.107 |
| 0.036 | 0.083 | 0.032 | 0.090 | 0.042 | 0.101 |
| 0.036 | 0.077 | 0.044 | 0.093 | 0.042 | 0.089 |
| **0.037** | **0.081** | **0.039** | **0.092** | **0.043** | **0.099** |
| 2.20 | | 2.36 | | 2.32 | |

**Fig. S12** Target engagement study in treated and untreated dogs
